# ViraLM: Empowering Virus Discovery through the Genome Foundation Model

**DOI:** 10.1101/2024.01.30.577935

**Authors:** Cheng Peng, Jiayu Shang, Jiaojiao Guan, Donglin Wang, Yanni Sun

## Abstract

**Motivation:** Viruses, with their ubiquitous presence and high diversity, play pivotal roles in ecological systems and have significant implications for public health. Accurately identifying these viruses in various ecosystems is essential for comprehending their variety and assessing their ecological influence. Metagenomic sequencing has become a major strategy to survey the viruses in various ecosystems. However, accurate and comprehensive virus detection in metagenomic data remains difficult. Limited reference sequences prevent alignment-based methods from identifying novel viruses. Machine learningbased tools are more promising in novel virus detection but often miss short viral contigs, which are abundant in typical metagenomic data. The inconsistency in virus search results produced by available tools further highlights the urgent need for a more robust tool for virus identification.

**Results:** In this work, we develop a Viral Language Model, named ViraLM, to identify novel viral contigs in metagenomic data. By employing the latest genome foundation model as the backbone and training on a rigorously constructed dataset, the model is able to distinguish viruses from other organisms based on the learned genomic characteristics. We thoroughly tested ViraLM on multiple datasets and the experimental results show that ViraLM outperforms available tools in different scenarios. In particular, ViraLM improves the F1-score on short contigs by 22%.

**Availability:** The source code of ViraLM is available via: https://github.com/ChengPENG-wolf/ViraLM.

## 1 Introduction

Viruses are microscopic infectious agents that require living cells of organisms to survive and replicate. They are ubiquitous and affect every form of life, from bacteria to plants and animals [1, 2]. There are many types of viruses, which can be classified into different categories based on their genetic materials (RNA viruses vs DNA viruses), hosts (e.g., eukaryotic vs prokaryotic), morphology (e.g., tailed viruses), etc. Some eukaryotic viruses have significant implications for public health, such as causing infectious diseases or even pandemics. Given that numerous pathogens are zoonotic in nature, vigilant surveillance of viruses in animal populations and diverse ecosystems is a critical aspect of One Health initiatives. In addition, viruses that infect prokaryotes, such as bacteria and archaea, can regulate the abundance of those prokaryotes and thus can profoundly impact the composition and function of microbial communities. Due to the difficulties of isolating viruses in labs, metagenomic sequencing has become the major means to explore the diversity and functions of viruses within various ecosystems. For example, recent studies reveal the critical role of viruses in biogeochemical cycles within marine and terrestrial ecosystems [3, 4]. The studies on urban microbiomes suggest that urban microbiomes are ecologically distinct from those of soil and humans, representing distinct niches with different genetic profiles [5].

Despite important findings about viruses in many ecosystems, our understanding of the virome on Eartch is still limited. It is estimated that the total number of distinct virus species is about 10^7^ to 10^9^, far exceeding the number of viruses currently cataloged in the viral genome databases [6]. Thus, there are still a substantial number of undiscovered viruses on Earth. Recently, the advent of high-throughput sequencing enables researchers to directly sequence all the genetic materials from various samples, leading to a wealth of metagenomic sequencing data. Consequently, mining viral sequences from metagenomic data, which also encompasses genetic materials from bacteria, archaea, and even eukaryotes, has become a critical step in exploring the full diversity of viruses.

There are several challenges in identifying viruses from metagenomic data. First, metagenomic data contains reads of heterogeneous origins, such as bacteria, archaea, and eukaryotes. Different mechanisms such as horizontal gene transfer lead to shared regions between genomes of viruses and other organisms. For example, prophage is commonly identified in bacteria as a region shared by prokaryotic viruses and their hosts. Our empirical studies show that ∼76% viruses in the RefSeq database have significant alignments with their hosts [7]. Shared sequences between viruses and other organisms increase the difficulty in virus detection [8]. Second, viruses have high genetic diversity with a wide range of genome length and protein composition. The smallest bacteriophage genomes are around only 2,435 base pairs [2], while the sizes of nucleocytoplasmic large DNA viral genomes can exceed two million base pairs [9]. In addition to genome size, the protein composition of viruses also demonstrates a broad spectrum of diversity. Based on the previous study [10], many of the newly discovered viruses include novel genes without detectable homology to those in known viruses. In particular, while most RNA viruses have RNA-dependent RNA polymerase (RdRp) as a possible marker, many other types of viruses lack marker genes. Third, the current reference viruses in available databases are far from providing a complete representation of the vast diversity present in nature. As a result, alignment-based tools tend to miss novel viruses that do not possess significant similarities with the reference genomes. With these challenges, current tools often produce very different numbers of viral-like contigs in metagenomic data. Even for relatively well-characterized human gut samples, there is high variability in the percentage of recovered viruses (14-87%) [11], underscoring the need for a robust and accurate virus identification tool.

### 1.1 Related work

Several attempts have been made to tackle the virus identification task, most of which incorporate machine-learning algorithms in order to address the limitation of alignment-based methods [12, 13, 14, 15]. These methods can be roughly categorized into two types based on the main features they rely on: nucleotide-based methods and protein-based methods. Nucleotide-based methods directly leverage nucleotide sequences to distinguish viruses from other sequences. A representative tool in this group, DeepVirFinder [12], uses one-hot encoding to embed both the original sequence and its reverse complement. Then, the encoded sequences will be fed into a convolutional neural network (CNN) to learn the motif-related features automatically. While CNN has proven effective at discerning complex patterns within nucleotide sequences [16], its ability to capture long-range dependencies in DNA sequences is hampered by the need for predefined window sizes [17]. This intrinsic constraint may adversely impact the accuracy of virus detection by DeepVirFinder.

Some other virus detection tools use protein-based features so that they can leverage hallmark genes in viral genomes. For example, VIBRANT [13] annotates the predicted open reading frames according to the hidden Markov Model search results of protein families against the public databases. Then the resulting annotation metrics are analyzed by the fully connected neural network. VirSorter2 [14] employs a customized protein database containing viral hallmark genes manually collected from metagenomic data. Then, a series of random forest classifiers are trained on distinct viral groups to generalize their performance to diverse viral genomes. More recently, a hybrid method named geNomad [15] further improves the performance of protein-based methods by aggregating and incorporating the features from nucleotide sequences. geNomad employs a neural network to learn nucleotide-based information and a tree ensemble model to extract protein features. The scores from both models will be weighted to generate final predictions.

Despite the promising results, methods that rely on protein features typically require the presence of relatively complete protein organization in query sequences. This limitation restricts their ability to accurately predict sequences with few proteins or sequences derived from non-coding regions. For example, VIBRANT can not handle sequences that encode less than four proteins. Considering that the majority of viral contigs in metagenomic data are short [18], these methods often fail to recruit many viral contigs in the metagenomic data. In addition, some protein-based methods must use alignment-based output as features in the learning process. However, as new virus groups are discovered, the expansion of reference databases may eventually lead to a significant increase in computational resources and running time requirements for the alignment process. For example, VirSorter2 requires more than two hours to classify 5,000 contigs in our experiments.

### 1.2 Overview

To address the major challenges of the high diversity of viruses and the difficulty of distinguishing ambiguous genomic regions shared between viruses and other organisms, we introduce Viral Language Model (ViraLM), the first virus identification tool powered by a pre-trained general-purpose genome foundation model. Foundation models have achieved remarkable success in many domains such as natural language processing, computer vision, and graph learning [19, 20]. A prominent example is ChatGPT, utilizing the Generative Pre-trained Transformer (GPT)-3 [21] as its underlying foundation model. Foundation models are typically pre-trained on extensive and diverse datasets, facilitating their adaptability to specialized downstream tasks through fine-tuning. This pre-training, often employing self-supervised learning, enables the model to capture the important patterns and semantics in the training data. In our work, we employ the latest genome foundation model, DNABERT-2 [22] as the backbone of ViraLM. DNABERT-2 has been pre-trained on a vast array of organisms, acquiring valuable representations of DNA sequences, which is particularly useful for distinguishing viral sequences from those of other species.

To adapt the genome foundation model for virus detection, we fine-tune this model for a binary classification task with two labels: viral sequences vs. others. We constructed a substantial viral dataset comprising 49,929 high-quality viral genomes spanning diverse taxonomic groups as positive samples. To ensure that ViraLM can learn a precise decision boundary, we curated a negative training set by including sequences commonly misclassified as viruses, such as prokaryotic host genomes (bacteria and archaea) and eukaryotes that are frequent virus hosts, like insects. This strategic selection of challenging negative samples for the fine-tuning process enhances ViraLM’s learning efficacy and its applicability to complex real-world data. The inherent strengths of the foundation model, including strong learning ability, adaptability, and time and cost-effectiveness, combined with our specialized training on an extensive viral dataset, render our tool ViraLM both accurate and fast.

To our best knowledge, ViraLM is the pioneering tool for virus identification that harnesses a genome foundation model. We extensively tested ViraLM on several datasets, including the RefSeq benchmark dataset, the IMG/VR dataset, and three real metagenomic data. We benchmarked ViraLM with three leading protein-based tools (VirSorter2, VIBRANT, and geNomad) and one nucleotide-based tool (DeepVirFinder). Our experimental results show that ViraLM consistently outperforms these existing tools in various experiments, with a notable 22% improvement in the F1-score for short contigs.

## 2 Methods and materials

The framework of ViraLM is illustrated in Fig. 1. The main architecture of ViraLM contains two parts: a pre-trained Transformer block from DNABERT-2 and a fine-tuned binary classifier layer for virus classification. Utilizing the pre-trained DNABERT-2 as the backbone for ViraLM provides two main benefits. First, the self-supervised learning approach used in DNABERT-2 (Fig. 1 A) allows ViraLM to leverage genomic data without the need for laborious labeling. This overcomes the limitations of task-specific supervised learning, which relies on limited labeled datasets, and enables the model to capture more comprehensive motif-related information across diverse microbial genomes. Second, by initializing ViraLM with the knowledge (parameters) from DNABERT-2 (Fig. 1 B), the model can rapidly converge in the virus identification task through fine-tuning, resulting in accelerated training and prediction. In the prediction process (Fig. 1 C), the input sequence is automatically segmented into non-overlapping 2kbp fragments, which are then tokenized and fed into the Transformer block. Each fragment has a prediction score (between 0 and 1.0), indicating the likelihood of the input fragment being a viral sequence. The prediction scores of the segments from the same sequence are averaged to generate a final prediction.

**Fig. 1.**
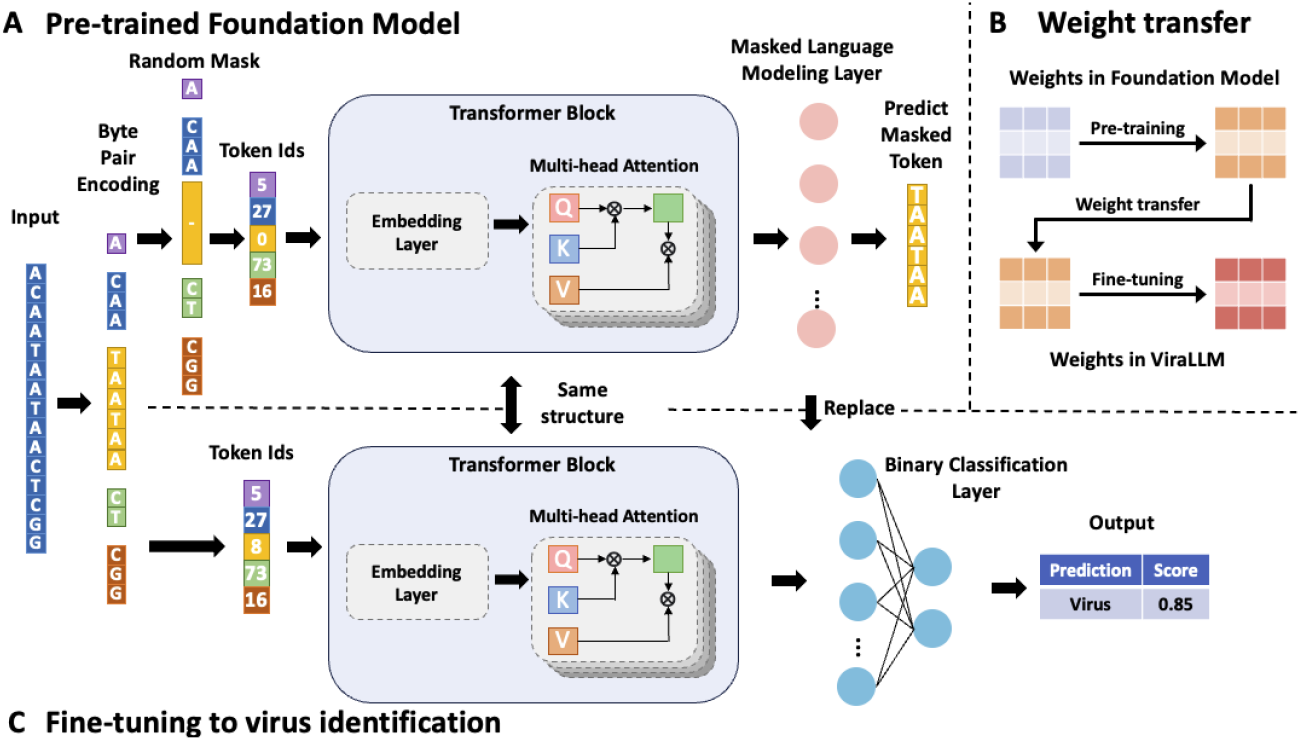
The framework of ViraLM. (A) The structure of the pre-trained foundation model DNABERT-2. Given an input sequence, it is first encoded into a token vector by Byte Pair Encoding. Then the tokens are randomly masked and fed into the Transformer black which contains an embedding layer and multi-head attention block. During pre-training, the model predicts the marked token based on the contextual information. (B) The pre-trained weights in the foundation model are transferred to ViraLM. (C) Then all the learnable parameters are fine-tuned together on the virus identification dataset. The input sequence is tokenized and fed into the transformer block. Then the binary classification layer will aggregate the result from the transformer block to generate a final prediction.

In the following section, we first introduce the Transformer block in DNABERT-2, which contributes significantly to the model’s powerful feature learning capacity. We will focus on the advancements it brings to enhance the efficiency and effectiveness of ViraLM. Then, we will elaborate on the modifications and adaptations made to this foundational model, specifically tailored to the virus identification task. Finally, we will outline the process of collecting and generating the datasets utilized in our experiments.

### 2.1 Basic structure of DNABERT-2

The overall structure of DNABERT-2 is shown in Fig. 1 A. To enhance the performance in genomic analysis, DNABERT-2 incorporates a series of advances and training strategies, including: (1) employing Byte-Pair Encoding (BPE) [23] to address the inefficiency of k-mer tokenization; (2) utilizing multi-head self-attention mechanism to capture contextual information; (3) pre-training on a multi-species dataset from 135 distinct species across seven categories. Below, we provide a concise overview of the three critical components that underpin the performance of the foundation model.

#### 2.1.1 Byte-Pair Encoding

In natural language processing, sentences can be divided into a list of words based on their semantics and grammar. However, when it comes to genomic sequences, the absence of naturally meaningful “words” poses a challenge for effective tokenization. Previous studies [24, 25] have employed k-mer tokenization, which involves splitting the sequence into overlapping substrings of a fixed length (k-mers). However, this approach faces information leakage issues because of the shared bases between adjacent k-mers. On the other hand, non-overlapping k-mer tokenization methods present a different challenge. Even a single mutation, insertion, or deletion in the sequence can substantially alter the tokenized embedding, making it difficult for the model to align similar sequences in a high-dimensional space [24].

Thus, Byte-Pair Encoding (BPE) is employed [23] to overcome the issues of k-mer tokenization. BPE is a statistical-based subword segmentation algorithm. Instead of using a pre-defined vocabulary of fixed-length k-mers, BPE dynamically learns a vocabulary of variable-length tokens based on the co-occurrence frequency in the datasets. During the tokenization process, BPE initially considers each nucleotide (A, C, G, and T) in the input sequence as an individual token. Then, BPE iteratively merges pairs or segments of nucleotides that commonly occur together in the genome. By leveraging the co-occurrence patterns of nucleotides, BPE dynamically constructs a vocabulary of variable-length tokens that captures meaningful subword units within the genomic sequences. This iterative merging process allows BPE to identify and encode biologically relevant information in the form of cohesive subword tokens, facilitating more effective representation and analysis of genomic data.

With the vocabulary generated by BPE, the input nucleotide sequences are translated into a sequence of tokens. An illustration of this process is provided in Fig. 1 C, where an input sequence is divided into five tokens of varying lengths: A, CAA, TAATAA, CT, and CGG. Then each token in the sequence is assigned a unique identifier known as a token ID. These sequences of token IDs serve as the input for the transformer block, allowing the model to effectively learn and capture the relationships and patterns within the genomic data.

#### 2.1.2 Transformer block

Because even short nucleotide sequences of 1kbp length can contain more than 230 BPE tokens, convolution and recurrence in CNNs and RNNs often struggle to capture these long-term token dependencies due to their fixed window size or vanishing gradients issue. To address this challenge, DNABERT-2 employs a 12-layer Transformer [17] to obtain contextual information within the sequences. The multi-head attention mechanism utilized in each layer allows access to all features in the sequence regardless of token distance or order.

In the transformer block, the tokenized sequence is first encoded into a matrix representation, denoted as *M* ∈ *ℝ*^L*×*embed^. Here, each token is mapped to a high-dimensional embedding space, where L represents the length of the tokenized sequence, and embed corresponds to the dimensionality of the embedding space. This encoding process ensures that tokens with semantic relationships are positioned closely together in the high-dimensional embedding space, enabling effective representation of the contextual and semantic information between different tokens. Then, the matrix is fed into the multi-head self-attention layer that can be expressed as Eqn. 1.

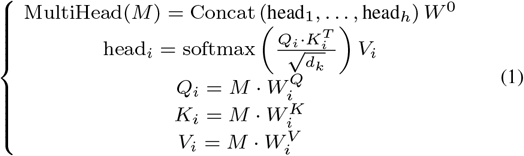

Where the input matrix *M* is projected into queries (*Q*), keys (*K*), and values (*V*) in dimension *d*_*k*_ for each attention head *head*_*i*_ separately. *W* ^0^, 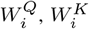 and 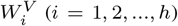 are learned matrices of the linear projection. Here, the query represents the information being sought, while the key serves as a reference for comparison. By computing attention scores between query-key pairs, the model determines the relationship between tokens. These attention scores are then used to weigh the corresponding values. By projecting the embedded matrix into separate queries, keys, and values for each attention head, the model can capture diverse perspectives and learn different representations of the sequence. This mechanism enables the model to assign distinct attention weights to individual tokens based on their relevance to one another, effectively leveraging the relationships between the sequence motifs in the sequence.

#### 2.1.3 Pre-training on multi-species genomes

Recent research has shown that machine learning models trained on diverse, multi-species genomic data tend to outperform those trained on single-species data [25]. This underscores the importance of using a large and diverse dataset to enhance a model’s learning capacity. However, the existing methods often suffer from the limited number of high-quality labeled data, which can hinder their ability to effectively extract most informative patterns.

To address the limitation, DNABERT-2 is pre-trained on an expansive dataset containing 32.5 billion nucleotide bases, sourced from the genomes of 135 different species across seven diverse categories, including bacteria, fungi, protozoa, mammals, and other vertebrates. This pre-training utilizes the Masked Language Modeling (MLM) technique [26], a self-supervised learning strategy that enables the model to discern general patterns and motifs from the unlabeled data, independent of any specific task. During pre-training, approximately 15% of the input tokens are randomly selected and replaced (masked) with either a specific [*MASK*] token or a randomly chosen different token. Then the model is tasked with predicting these masked tokens based on their surrounding context. By employing MLM, DNABERT-2 gains the ability to comprehend the various contexts in which each nucleotide base appears, thus enabling a comprehensive understanding of genomic sequences. As a result, DNABERT-2 effectively leverages the vast amount of genomic data obtained through metagenomic sequencing.

### 2.2 Adapt DNABERT-2 to virus identification task

To adapt the foundation model for the virus identification task, a fine-tuning method is applied, combining the understanding of genomic sequences obtained during the pre-training phase with the task-specific dataset. This approach leverages the strengths of both self-supervised and supervised learning, enabling the model to gain a comprehensive understanding of genomic patterns and contexts while being specifically tailored to virus identification.

First, we create a new model with a structure similar to DNABERT-2 but designed specifically to classify input sequences as virus or non-virus. To accomplish this, a fully connected layer is integrated behind DNABERT-2’s attention layer to act as a binary classifier during prediction. The learnable parameters of this new model are initialized by transferring values from the corresponding parameters in DNABERT-2’s attention block (see Fig. 1 B), ensuring that the model starts with the parameters and representations learned by DNABERT-2. Second, unlike DNABERT-2’s self-supervised training, ViraLM will be trained using labeled data, which consists of positive samples (virus sequences) and negative samples (non-virus sequences) generated from high-quality genomes. During fine-tuning, the learnable parameters are aligned with the specific training objective of virus identification. To facilitate parameter updates, binary cross-entropy loss and the Adam optimizer with a learning rate of 3e-5 are employed. Once the fine-tuning process is completed, the model becomes capable of accurately identifying virus sequences from the input data.

### 2.3 Data collection and experimental setup

#### 2.3.1 Datasets

We rigorously tested ViraLM on multiple datasets with increasing complexity. The main information of the test data is listed in Table 1 And we will detail how we construct these datasets in the following sections.

**Table 1.** Summary of the datasets used in the experiments.

| Dataset | Description | Type | Number of contigs |
| --- | --- | --- | --- |
| Fine-tuning data |  |  |  |
| Training data | The positive data (49,929 viral genomes) are complete viral genomes downloaded from the NCBI RefSeq and high-quality viral genomes collected by VirSorter2 [14]. The negative data (245,734 non-viral sequences) are complete assemblies of prokaryotes (bacteria and archaea), eukaryotes (fungi and protozoa), and chromosomes downloaded from the NCBI RefSeq. The genomes within each group (positive and negative) are randomly divided into training and test sets with a ratio of nine to one. Then, we randomly cut the genomes into short contigs ranging from 300bp to 2kbp to simulate variable-length contigs in the metagenomic data. | Randomly cut contigs (300bp to 2kbp) from training genomes. Balanced the number of positive and negative contigs. | 36,617,141 |
| Test data | The gained test genomes are filtered to exclude highly similar sequences to the training data with a stringent threshold of 50% overall coverage. The test set also includes genomes of bats and humans downloaded from NCBI to evaluate the model’s generalization to other host organisms. All the genomes are fragmented into short contigs and sampled to balance positive and negative samples. | Randomly cut contigs (500bp to 20kbp) from test genomes. Balanced the number of positive and negative contigs. | 434,443 |
| Diverse viruses data |  |  |  |
| IMG/VR v4 [27] | A total of 367,782 genomes, representing a broad range of novel viruses, are collected from the IMG/VR v4 database. We cut the genomes into 2kbp length fragments with 0 (non-coding regions), 1 and 2 proteins, and 5kbp length fragments with 3, 4, and 5 proteins to generate the test dataset. | Database | 908,116 |
| Real sequencing data |  |  |  |
| Water [28] | Metagenomic data sequenced from surficial water of Jinghu Lake (China) | Assembled contigs | 1,294,985 |
| Soil [29] | Metagenomic data sequenced from *Phragmites* plants soil | Assembled contigs | 461,310 |
| Insect [30] | RNA-Seq data sequenced from sandflies | Assembled contigs | 149,322 |

*Labeled data for fine-tuning* To construct a task-specific training dataset for fine-tuning, we integrated two virus datasets from different sources to generate positive samples. First, we downloaded the complete viral genomes released before September 2023 from the National Center for Biotechnology Information (NCBI) RefSeq database (https://www.ncbi.nlm.nih.gov/). The RefSeq viral genomes are divided based on their realm-rank taxons: *Adnaviria, Duplodnaviria, Monodnaviria, Riboviria, Ribozyviria* and *Varidnaviria*. Specifically, we combine *Adnaviria, Ribozyviria*, and genomes that lack realm-rank taxons into one group due to their small sizes. Second, we downloaded a dataset of high-quality viral genomes used in VirSorter2 [14], which were carefully collected and curated in the previous works. These genomes represent a set of novel viruses that are dissimilar to RefSeq reference genomes. They are pre-grouped into dsDNA phages, *Lavidaviridae* (virophages), NCLDVs (giant viruses), and RNA viruses. In total, the virus dataset contains 49,929 genomes.

When considering negative samples for training, it is impractical to include all possible non-viral genomes. Instead, a rigorous sampling approach was employed, focusing on sequences that are prone to being misclassified as viruses. These include the host genomes of prokaryotic viruses, such as bacteria and archaea, as well as eukaryotes that are known to commonly carry viruses, such as insects. By incorporating these challenging cases, the model can learn a more effective decision boundary between viruses and non-viral entities. First, we downloaded complete assemblies of prokaryotes (bacteria and archaea) and eukaryotes (fungi and protozoa) from the NCBI RefSeq database. These carefully selected genomes represent typical non-viral entities commonly appearing in metagenomic data [30, 31]. Additionally, we also include the plasmid data from the RefSeq database, which are prone to be misidentified by the existing virus identification tools. Furthermore, considering that insects serve as hosts and key vectors for carrying a wide range of viruses, including arthropod-borne viruses (arboviruses) that pose significant public health concerns [32], we also include the chromosomes from the *Insecta*. Due to the significantly larger size of the bacteria and *Insecta* datasets compared to the other non-viral organisms, we down-sampled two genomes per genus from the bacteria datasets and one genome per genus from the *Insecta* dataset. To remove the possible endogenous virus elements in non-viral sequences that might confuse the model, we apply BLASTN [33] to align the non-viral sequences against the viral sequences and remove the common regions from the sequences.

Finally, the genomes within each group (positive and negative) are randomly divided into training and test sets with a ratio of nine to one. This separation was performed individually for each realm-rank taxon to ensure a balanced distribution of different organisms in the datasets. The genomes are randomly cut into short contigs ranging from 300 to 2kbp to simulate variable-length contigs in the metagenomic data. To mitigate data imbalance caused by significant differences in genome sizes, the fragments from the non-viral genomes are down-sampled to match the size of the virus dataset.As a result, we construct an extensive training set consisting of 40 billion nucleotide bases from bacteria, archaea, plasmid, fungi, protozoa, *Insecta*, and viruses. The large volume and diverse composition of the training data enable the model to gather ample information, enhancing its ability to generalize effectively when processing previously unseen species.

To ensure a fair evaluation of the model, we examine the sequences in the test set to exclude those that exhibit high similarity with the training sequences. This is achieved by aligning the test sequences using BLASTN to the training data. Alignments from different regions of the same query sequence are aggregated to calculate the overall alignment coverage and identity. According to the guidelines provided by the International Committee on Taxonomy of Viruses (ICTV) [34], a similarity threshold of 70% is established for classification at the genus level. To ensure a rigorous testing process, we adopt a more stringent threshold, requiring an overall coverage of less than 50% between training and test sequences. Thus, the test set can provide a fair and unbiased assessment of the model’s performance. In addition, we include in our test set the reference bat genome (order *Chiroptera*) and human genome to further test the model’s generalization ability on other large eukaryotic organisms that are not included in our training data. Similar to previous steps, these sequences are also fragmented into short contigs (500bp to 20kbp). In particular, all the test sequences are sampled to balance the positive samples and negative samples, ensuring a fair evaluation.

*IMG/VR dataset* The Integrated Microbial Genome/Virus (IMG/VR) v4 database [27] contains 5,576,197 high-confidence viral genomes collected from different environment samples. These genomes are systematically quality-checked and classified into high-quality, medium-quality, low-quality, and unsure-quality genomes by IMG/VR. We recruit 495,576 high-quality genomes from the database. These genomes are aligned against the training set to remove highly similar sequences with BLASTN (i.e., overall coverage >50%). After the removal process, we get a total of 367,782 genomes, which represent a broad range of novel uncultured viruses. These diverse viruses may exhibit various protein densities. To test how the number of proteins on a contig can affect the performance of virus identification, we first predict the open reading frames using Prodigal [35]. Then the genomes are cut into 2kbp length fragments with 0 (non-coding regions), 1 and 2 proteins, and 5kbp length fragments with 3, 4, and 5 proteins, respectively.

*Real sequencing dataset* We retrieved two public real metagenomic data and one RNA-Seq data sampled from different environments, including water [28], soil [29], and insect-associated ecosystems [30], respectively. The water metagenome is sequenced from a total of 30 surficial water samples collected from Jinghu Lake (China), a lake that is replenished by reclaimed water. The soil microbiome comes from *Phragmites* plants rhizosphere soil (soil that is firmly attached to root) and bulk soil (nearby soil without *Phragmites* plants). The insect microbiome is obtained from hundreds of sandflies. Low-quality reads and adaptor sequences are removed from the raw sequencing data using Fastp [36]. Then the resulting clean reads are assembled into contigs using Megahit [37]. Only the contigs with lengths over 2kbp are reserved for prediction, resulting in 1,294,985, 461,310, and 149,322 contigs in the three types of samples, respectively. Table 1 summarizes the key information of these real sequencing datasets.

#### 2.3.2 Metrics

Following previous works, we employ precision, recall, and F1-score as metrics for evaluating virus identification performance. The formulas are listed below (Eqn.2, Eqn. 3, And Eqn. 4):

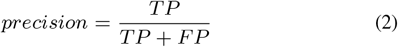

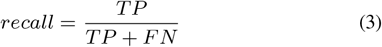

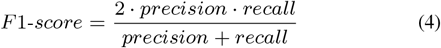

True positive (TP), false negative (FN), and false positive (FP) represent the number of corrected identified viruses, the number of viruses misclassified into non-viruses, and the number of falsely identified viruses, respectively.

## 3 Result

In this section, we evaluate ViraLM on various test datasets and compare it against new or state-of-the-art methods including geNomad [15], VirSorter2 [14], VIBRANT [13] and DeepVirFinder [12]. Because VirSorter2, VIBRANT, and geNomad rely on their curated category-specific protein databases for prediction, we directly run these tools without retraining. Although DeepVirFinder supports retraining, it can not process a large-scale dataset because of the limitation of one-hot encoding. We run VirSorter2 (v2.2.4) with the option “–include-groups dsDNAphage,NCLDV,RNA,ssDNA,lavidaviridae” to predict viruses from all the groups. VIBRANT v1.2.1, geNomad v1.7.4, and DeepVirFinder v1.0 are run on their default settings. Similarly, during all the following tests, ViraLM is run under its default setting without any extra adjustment for different test datasets.

### 3.1 Ablation study in ViraLM

As ViraLM is built upon a pre-trained foundation model, we first investigate whether the fine-tuning process, which adjusts the DNABERT-2’s parameters based on our virus identification task, brings any benefits, compared with starting from random parameters in the Transformer block. Specifically, we compare the performance of ViraLM using two different weight initialization strategies: random initialization and initialization with the pre-trained foundation model. For the randomly-initialized setting, we construct a new model with the same structure as ViraLM. In this model, all the learnable weights in the transformer block are randomly initialized using a normal distribution with a mean of zero and a standard deviation of 0.02. The biases in the model are initialized to zero. During the training process, we keep track of the training losses and F1-score for both models on the test set in each epoch. As shown in Fig. 2, the pre-trained model demonstrates faster convergence compared to the model with random initialization, resulting in superior performance with the same number of epochs. The observed trends highlight the importance of the foundation model as a valuable starting point for improving model accuracy and speeding up the training process. We also compare their final performance on contigs with various lengths in Fig. 3. The results show that the pre-trained model outperforms the model with random initialization across all contig ranges, underscoring the effectiveness of the foundational model in boosting overall performance.

**Fig. 2.**
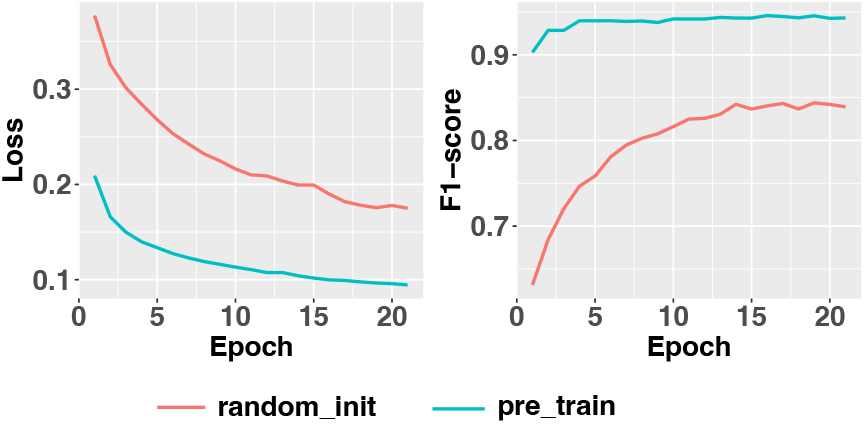
Comparison of training loss and F1-score between the model initialized using the pre-trained foundation model and the model with randomly initialized weights. The model initialized using the pre-trained foundation model converges faster and performs better.

**Fig. 3.**
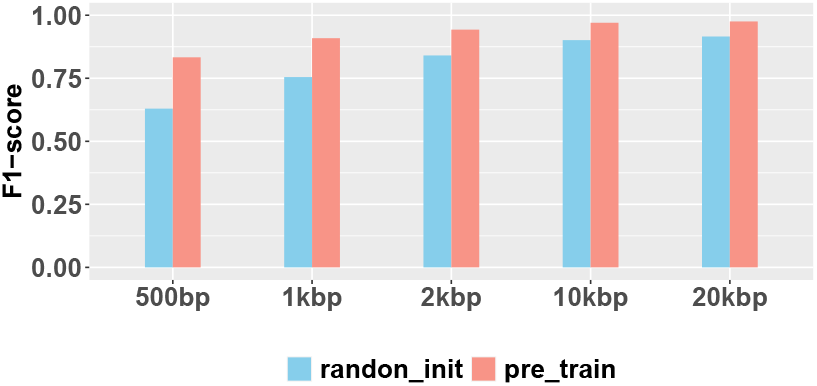
Comparison of virus identification performance on contigs of various lengths between the model initialized using the pre-trained foundation model and the model with randomly initialized weights.

### 3.2 ViraLM outperforms the benchmarked methods

We compared ViraLM with the state-of-the-art or new tools on the test set consisting of virus sequences from NCBI RefSeq and uncultured viral genomes obtained from metagenomes as detailed in Table 1. In order to evaluate whether prokaryotic or eukaryotic sequences pose a bigger challenge for virus detection, we evaluate and report the performance of all tools on the two types of negative test data separately.

*Prokaryotic genomes* First, we report the performance on the negative test set containing sequences from bacteria, archaea, and plasmids. Still, to ensure a balanced test set, we choose equal number of negative sequences as the viral sequences. After obtaining the prediction results, we draw an ROC curve using the prediction scores of each tool in Fig. 4. The area under the ROC curve reveals that ViraLM returns more reliable results compared with other tools. While Fig. 4 is generated on all the contigs in the test set, we further analyzed the virus identification performance of different tools on contigs of different length ranges. In general, the shorter the contig, the harder to distinguish viruses from others. As generating ROC curves for all length ranges are too tedious, we show the F1 score of all tools under their default setting. The results can be found in Fig. 5.

**Fig. 4.**
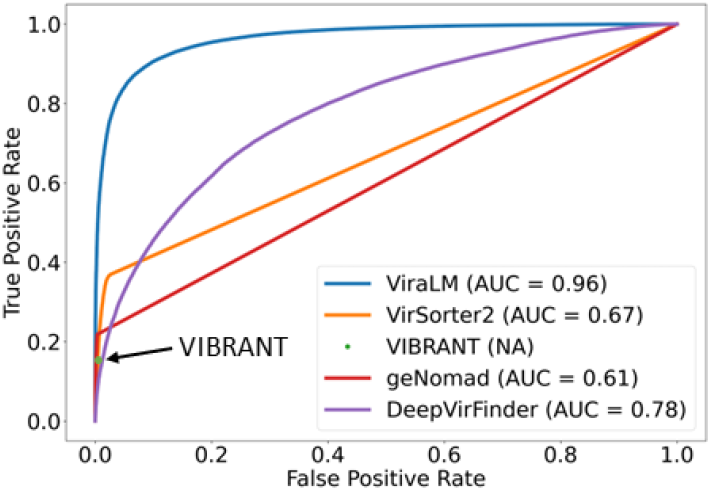
ROC curves on the contigs where negative samples only consist of prokaryotes (bacteria, archaea, plasmid). The value in the parentheses represents the AUC for each tool. Because VIBRANT does not provide a score associated with each prediction, we only plot one data point on the graph.

**Fig. 5.**
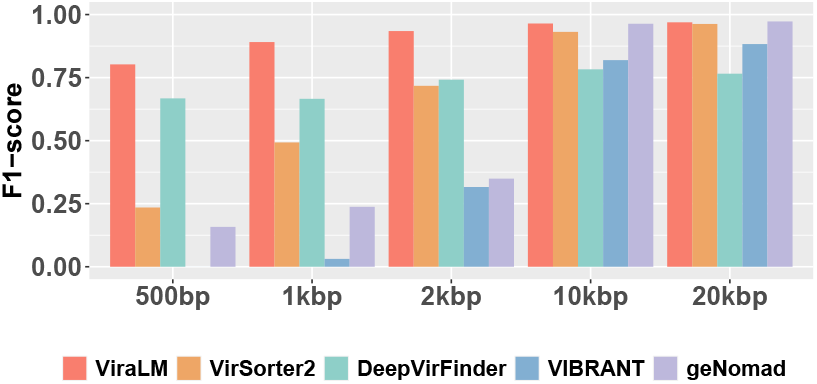
The performance of each tool on various-length contigs where negative samples only consist of prokaryotes (bacteria, archaea, plasmid). All tools are run under their default setting and prediction score cutoffs.

All the tools are run on contigs with lengths 500bp, 1kbp, 2kbp, 10kbp, and 20kbp, respectively. The comparison reveals that ViraLM outperforms other methods across all length ranges, highlighting its robustness for virus identification. The performance of all tools improves as the contig length increases, likely because of more available information carried in longer sequences. Specifically, all the protein-based methods (VirSorter2, VIBRANT, and geNomad) perform well on contigs over 10kbp. However, they tend to struggle with short contigs due to their reliance on complete protein information, which is less likely to be present in shorter sequences. VIBRANT can not predict on contigs shorter than 1kbp and therefore its score is zero. DeepVirFinder performs the second best on contigs under 2kbp, highlighting the strength of nucleotide-based methods. Yet its performance shows only a slight improvement, approximately 0.04, when contig lengths increase from 2kbp to 10kbp. This modest improvement is attributed to the inherent limitations of CNNs in leveraging long-term dependencies, compared to the more adept Transformer model.

*Eukaryotic genomes* Next, to gain a more comprehensive evaluation of ViraLM, we follow the design in [14] and extend our assessment to eukaryotic genomes, broadening the scope beyond small eukaryotes like fungi and protozoa. We conduct a comparative analysis of all methods on contigs of mixed length where negative test samples only contain contigs from five distinct eukaryotic groups (fungi, protozoa, insects, bats, and humans), which are frequently found in host-associated microbiomes. To assess the performance of each tool across various eukaryotic groups, we run each tool separately on these groups. The initial observation indicates that the ability of all tools to differentiate viral sequences from those of prokaryotic or eukaryotic origin is remarkably consistent, as evidenced by Fig. S1 in the supplementary file. This suggests that distinguishing viruses from these two categories of non-viral data has a comparable level of difficulty. A more detailed comparison of the tools’ precision, recall, and F1-score for each eukaryotic group is presented in Fig. 6.

**Fig. 6.**
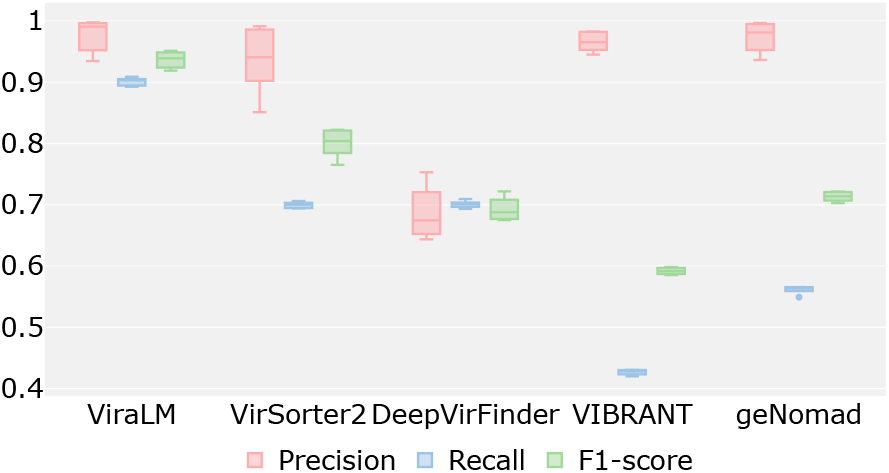
The performance of each tool on distinguishing viruses from every group of eukaryotic contigs: fungi, protozoa, insects, bats, and human. ViraLM outperforms the other tools across precision, recall, and F1-score. VirSoter2, VIBRANT, and geNomad exhibit high precision but show low recall.

Upon assessing the classification results, we find that ViraLM delivers an enhanced overall performance compared to the other methods. In general, ViraLM, VirSoter2, VIBRANT, and geNomad demonstrate high precision in all eukaryotic groups. The well-constructed viral hallmark gene databases contribute to the high precision of protein-based methods. However, VirSoter2, VIBRANT, and geNomad exhibit low recall, indicating that they are conservative models that are prone to generate fewer false positives but will miss a considerable number of true positives. In contrast, ViraLM achieves comparable precision to that of the protein-based tools while maintaining high recall across all groups. In particular, the F1-score of distinguishing viruses from bat and human-originated contigs are 0.91 and 0.92, respectively. As the closest species in the labeled training data for ViraLM is merely *Insecta*, achieving such high F1-scores underscores ViraLM’s adaptability and efficacy in analyzing metagenomic data derived from various and new host organisms.

### 3.3 Performance on IMG/VR dataset

We further benchmarked all the methods on the IMG/VR dataset [27]. Recently, IMG/VR presented its fourth version of the database, consisting of over 15 million viral genomes and genome fragments. The viruses in the database are systematically identified from a broad range of environmental samples and have undergone rigorous quality checks, annotations, and taxonomic classification. Thus, such an abundant and diverse virus dataset serves as a good third-party test set for evaluating the sensitivity of various virus identification methods. We recruit 367,782 high-confidence, high-quality genomes from the dataset. To evaluate the impact of protein number on the virus identification can affect the performance, we first predict the encoded proteins using Prodigal [35], then we cut the genomes into fixed-length (2kbp and 5kbp) contigs containing various number (zero to five) of complete proteins. Specifically, contigs that do not contain proteins indicate the non-coding regions in the genomes. Since there are only virus sequences in the experiments, we report the percentage of identified viruses of each tool in Fig. 7.

**Fig. 7.**
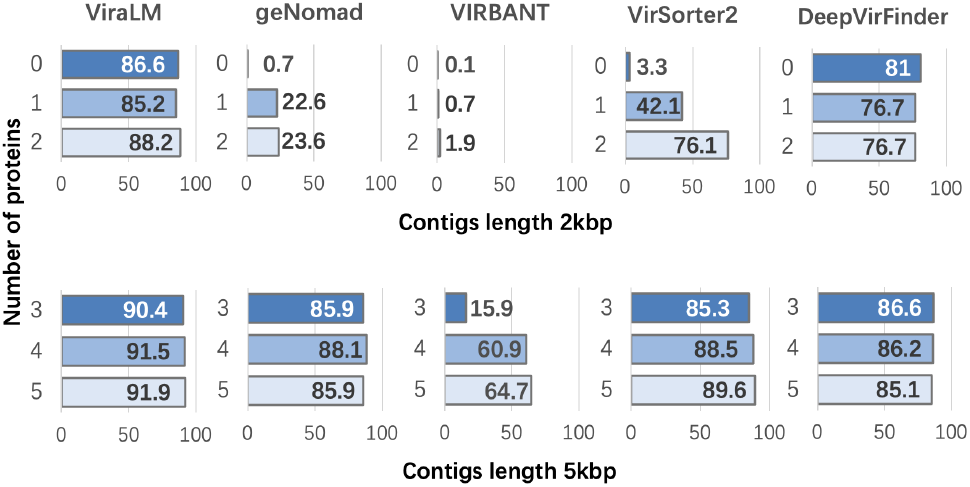
Sensitivity of virus identification on contigs with various protein densities, grouped by contig lengths. Top: 2kbp contigs; bottom: 5kbp contigs. Columns represent the compared tools. In each panel, the X-axis is the percentage of identified viruses (sensitivity); the Y-axis is the number of proteins encoded on the contigs.

The results reveal that ViraLM and DeepVirFinder identify most of the viral contigs across different protein densities. In particular, ViraLM and DeepVirFinder can accurately predict the contigs from non-coding regions, whereas protein-based tools commonly fail. Notably, ViraLM’s performance remains relatively stable across contigs of identical lengths, demonstrating that ViraLM can effectively utilize the nucleotide information. In contrast, the increased number of proteins leads to a significant increase in recall of geNomad, VIRBANT, and VirSorter2. This observation is consistent with the fact that they rely on marker proteins for prediction. In addition, the results show that geNomad and VIRBANT require more proteins for prediction than VirSorter2. Overall, the comparison across various contig lengths and protein densities highlights ViraLM as a reliable tool for uncovering the extensive diversity of viruses.

### 3.4 Performance on real sequencing data

Finally, we run all the methods on two public real metagenomic datasets and one RNA-Seq dataset released in previous studies [28, 29, 30]. These datasets, derived from distinct ecosystems such as aquatic environments, terrestrial soils, and insect populations, represent a wide range of microbial communities. In total, there are 1,294,985, 461,310, and 149,322 contigs with lengths over 2kbp, respectively. The number of viruses predicted by each tool and the overlap (intersection) among different groups is displayed by UpSet Plots in Fig. 8. Across all the experiments, DeepVirFinder always returns the largest number of predicted viral contigs. As shown in Fig. 6, DeepVirFinder has lower precision than other tools, suggesting that some of its returned predictions are not likely viruses. On the contrary, VIRBANT always returns the smallest number of predictions, which is also consistent with the result in Fig. 6. Fig. 8 exhibits a unanimous identification of 70,496 viruses in the water sample made by all tools, whereas in the insect microbiome, the tools’ predictions have small overlaps. This observation reveals that different ecosystems have different virus composition and complexity.

**Fig. 8.**
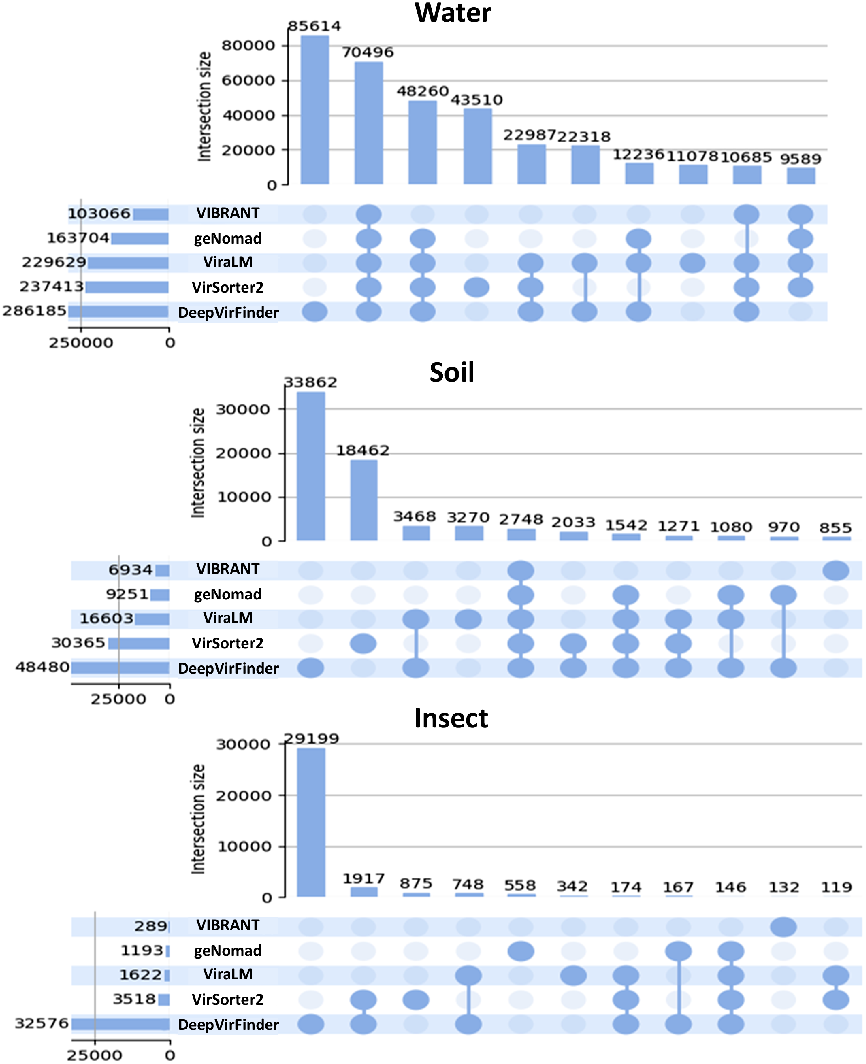
Comparison of the overlap of predicted viruses from different environmental samples. Each figure represents one of the three real datasets. In the figure, horizontal bars represent the number of viruses predicted by each tool. Each column represents a set of viruses. The vertical bar on top of the column shows the size of the set. The filled-in cells joined by a line indicate which groups share these viruses. The bars are sorted by the size of the intersection. For example, in the first figure, 70,496 viruses are identified by all the tools, while 11,078 viruses are uniquely discovered by ViraLM.

Although we lack the ground truth about the viral composition in these real data, we can decide the labels of some contigs based on very strict alignment results against reference genomes. We categorize the contigs into bacteria, archaea, eukaryotes, and viruses by assigning labels according to the best-hit BLASTN alignment against the NT database with a strict threshold (identity >85% and coverage >85%). As a result, 21,168, 31,119, and 1,292 contigs are classified into one of these groups. By summarizing the BLASTN results and analyzing the composition of the samples, we found that non-virus contigs outnumber virus contigs by approximately 28 to 316 times. Such pronounced imbalance can inherently impact the precision calculations across all methods. The performance of all the tools on these labeled subsets is shown in Fig. 9. The result confirms that DeepVirFinder tends to overestimate the virus population, resulting in the lowest precision. VIBRANT maintains relatively high precision and F1-score, while its recall varies much within different samples. In contrast, VirSorter2 and geNomad exhibit wide ranges of precision, varying from 0.05 to 0.88, demonstrating their limitation on specific environmental samples. In general, ViraLM presents the highest and most stable performance in real sequencing data sampled from different environments. As mentioned earlier, as the number of non-viral contigs is significantly larger than viral contigs, precision is naturally lower even when the false positive (FP) rate is low. For example, the FP rate of ViraLM in the water sample is 0.011, indicating that only 1% of the non-viral contigs are misclassified as viruses. However, given the large total number of non-viral contigs, a low FP rate can still contribute to a low precision calculation.

**Fig. 9.**
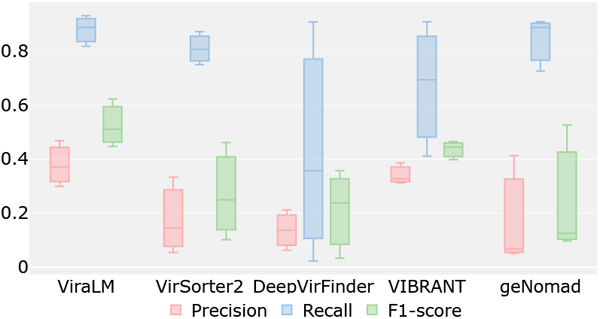
The performance on three real sequencing data by each tool. X-axis: the names of the tools. Y-axis: the value of the metrics on three different samples (represented by boxplots for easy comparison).

#### 3.4.1 Time efficiency and disk size

Because VirSorter2, DeepVirFinder, VIBRANT, and geNomad do not support running on GPU, we run these methods on Intel^®^ Xeon^®^ Gold 6258R CPU with 8 cores. In contrast, ViraLM is run on a single NVIDIA GeForce RTX 3090 GPU. All the methods are run on a test set containing 5,000 contigs. The elapsed time is shown in Table 2. Additionally, we record the sizes of disk space needed by the tools, predominantly comprising the databases and models. The rapid prediction capability and relatively compact size suggest that ViraLM is a lightweight and efficient tool that may assist and accelerate the discovery of new viruses.

**Table 2.** The elapsed time to make predictions on the test set and the size of disk space needed by the tools.

| Program | ViraLM | VirSorter2 | DeepVirFinder | VIBRANT | geNomad |
| --- | --- | --- | --- | --- | --- |
| Elapsed time(min) | 3 | 156 | 11 | 38 | 21 |
| Disk size(MB) | 447 | 12288 | 114 | 11264 | 1433 |

## 4 Discussion

In this work, we present a novel language model, named ViraLM, for virus identification. The major improvement of our method stems from the adoption of the pre-trained foundation model and our careful construction of the extensive training set. The pre-trained foundation model provides an extensive knowledge of the intricate patterns and relationships within genomes. Built on this prior knowledge, our model is able to recognize the subtle differences hidden in the nucleotide sequences between viruses and other organisms, leading to improved performance. In addition, we construct a vast dataset with novel uncultured viruses and distinct groups of non-virus organisms. This inclusion of additional genomes expands the diversity of our training data, enhancing the model’s generalization capability. Through benchmark experiments conducted on standard benchmark datasets, IMG/VR datasets, and real metagenomic data, our model consistently outperforms existing methods across various scenarios. In particular, it achieves a remarkable 22% improvement in F1-score on short contigs. Moreover, our model exhibits superior robustness when confronted with complex real metagenomic samples, highlighting its adaptability and advanced performance in different environments.

Although ViraLM has greatly improved virus identification, there are still some limitations. Firstly, although language models have the ability to capture long-range interactions in nucleotide sequences, they often encounter significant computational challenges when dealing with lengthy input sequences. This is particularly relevant considering that nucleotide sequences can reach lengths of millions of base pairs (bp). Second, due to the specific design of the training process, ViraLM may encounter limitations in detecting pro-virus fragments that are integrated within host genomes. Thus, we have several goals to optimize or extend ViraLM in our future work. First, we will investigate whether some length-invariant features or data compression methods can be incorporated into ViraLM to overcome the length limitation. Second, we will actively keep pace with the advancements in deep learning to maintain the computational efficiency of ViraLM. Third, we will explore the incorporation of additional features, such as binding sites and methylation sites, into our framework. By leveraging these supplementary features, we seek to enhance the sensitivity of ViraLM in identifying pro-virus fragments.

## Data Availability

All data and codes used for this study are available online via: https://github.com/ChengPENG-wolf/ViraLM.

